# Local Gazetteers Reveal Contrasting Patterns of Historical Distribution Changes between Apex Predators and Mesopredators in Eastern China

**DOI:** 10.1101/2020.07.15.202390

**Authors:** Kaijin Hu

## Abstract

**Background:** Humans have been causing the sixth wave of mass extinction of biodiversity. The situation of predators, especially of carnivores, is a key indicator of biodiversity, and the mesopredator release is a typical phenomenon in ecosystem recess. Local gazetteers are a rich resource for historical biodiversity research. However there are obvious biases in previous studies focusing only on presence records and neglecting the absence records. I recollected and analyzed the records by fixed methods to research historical biodiversity change.

**Methods:** Innovatively, this research used both presence and absence records from local gazetteers to reconstruct the distribution of 8 kinds of mammalian predators (i.e. large carnivores: tigers, leopards, and bears; medium-sized carnivores: wolves, foxes, civets, dholes, and mustelids) in eastern China from 1573C.E.(Common Era) to 1949C.E. (sorted into 4 periods). Then by utilizing statistics methods, e.g. Partial Least Squares Structural Equation Modeling(PLS-SEM), I analyzed the distribution changes(overall and sorted into 5 altitude groups), and the relation between carnivores and the influence from humans.

**Results & Conclusions:** I reconstructed the specific distribution changes of the carnivores and found the changes match the mesopredator release phenomenon. Furthermore, I found the specific relation(e.g. trade-off effect) between carnivores and the influence of humans. Besides, I found that bears and civets may be potential members of the “Tigers - Leopards - Dholes” predator guild.

**Prospects:** I have also collected records for other wild animals from local gazetteers in China. Based on this collection, I have built the Database of Wild Mammal Records in Chinese Local Gazetteers and am building the Database of Wild Bird Records in Chinese Local Gazetteers. I aim to continue relevant studies using these databases in the future.

## 1. Introduction

It has been widely accepted that Earth is facing a biodiversity crisis (Rahbek & Colwell 2011). In the past 500 years, a wave of extinction and declines in local animal population have been recorded, with both the rate and magnitude of this recent period of extinction being comparable to the five previous periods of mass extinction (Pimm *et al*. 1995; He & Hubbell 2011; Dirzo *et al*. 2014). Unlike these five previous periods of mass extinction which were primarily driven by global changes in climate and atmospheric-ocean chemistry, volcanism, and bolide impacts (Barnosky *et al*. 2011), the modern loss of biodiversity is considered as the consequence of human socioeconomic activities (Dirzo & Raven 2003; Ceballos *et al*. 2015). Humans have been causing the sixth wave of extinction through changing global climate, overkilling species, overexploitation resources, introducing exotic species, spreading infectious diseases, and destructing and modifying habitats (Leakey & Roger Lewin 1996; MacPhee *et al*. 1999; Dirzo & Raven 2003; Thomas *et al*. 2004; Martin 2005; Jackson 2008; Wake & Vredenburg 2008; Ceballos *et al*. 2015; Yap *et al*. 2015).

Although there is no doubt that the impact of the recent “sixth period of mass extinction” has extended across distinctive taxonomic groups, each species responds differently to climatic shifts, habitat fragmentation, and decline in species abundance caused by anthropogenic disturbance (Cardillo *et al*. 2008; Lorenzen *et al*. 2011). Moreover, the life history characteristics of different species are also to some extent correlated with their risk of being threatened by human activities (Davidson *et al*. 2009; Öckinger *et al*. 2010; Cardillo & Meijaard 2012). Despite this, such studies usually had poor predictive power to reliably link species’ vulnerability with their life history traits (Jablonski 2001, 2008; Pocock 2011). Even relatively well-established patterns of correlation, such as large body size being known to be related strongly to extinction risk in mammals, still failed to accurately predict the extinction risk of a specific species affected by human activities. This is due to (*i*) multiple ecological factors and human activities interact to affect species that differ by orders of magnitude in life history traits (Davidson *et al*. 2009), and (*ii*) human disturbance to specific species is usually regulated by the local community structures and population dynamics in which the species is embedded (Kitahara & Fujii 1994; Gill 2007; Tylianakis *et al*. 2007). A phenomenon termed “mesopredator release” represents one of the best-referred examples. Large-bodied predators were persecuted by humans, but the abundance of mesopredators might undergo dramatic increases as a result of trophic cascading effects in communities (Prugh *et al*. 2009; Wallach *et al*. 2015). Consequently, in view of the complex synergy of both intrinsic species traits and human’s role at different spatial-temporal and ecological scales, more specific quantitative assessments of the relationships between local extinction of endangered species and anthropogenic factors are urgently required (Fritz *et al*. 2009; Dirzo *et al*. 2014; Sandom *et al*. 2014; Wan *et al*. 2019; Teng *et al*. 2020).

In China, the distribution of wild animals has changed greatly over time, with many formerly widely distributed wild animals now endangered or even extinct (Chatterjee et al. 2012; Qian *et al*. 2020). Meanwhile, the population in China has increased significantly since the introduction of American crops in the 16^th^ century, which triggered prosperous socio-economic activities that may be responsible for the extinction of wild animals in China (Li 1998; Cao 2001). China has a long history of using local gazetteers to record momentous events and situations, including those related to local wildlife, and previous researchers have published numerous collection lists of animal records (He 1993; 1997; Wen 2009; 2013; 2018). By exploring these historical records in various ways, researchers have attempted to reconstruct the spatio-temporal distribution and extinction of engendered animals in China, and then disentangle the relative roles of anthropogenic and climate changes in shaping their distribution and extinction (Jiang *et al*. 1995; Li *et al*. 2002, 2020; Zhou & Zhang 2013; Li *et al*. 2015; Turvey *et al*. 2015, 2017; Wan & Zhang 2017; Nüchel *et al*. 2018; Zhao *et al*. 2018; Wan *et al*. 2019; Qian *et al*. 2020; Teng *et al*. 2020). However, such studies have relied on biased species observation data which only focused on the “presence” records of local animals but neglected the “absence” records. For each species, they often defined the time of extirpation as the last observation (or called last occurrence) time which is interpreted from the most recent record there.

The previous studies neglected the absence records and thus led to biases. The distribution of local gazetteers is not uniform in either space or time. If only considering the presence (1) records, it will greatly confuse the absence (0) areas and unknown(null) areas. The collection of data from gazetteer-rich and gazetteer-poor regions/periods may cause obvious biases in estimating the distribution of wild animals(Boakes *et al*. 2010). For example, there are few gazetteers in war-torn or sparsely populated areas, and there were only one or two gazetteers from many regions throughout history. Therefore, to accurately evaluate the spatio-temporal distribution and extinction of wild animals in China, it is necessary to take into account these biases in the data collected, and then do the re-collection job to track changes in biodiversity and analyze how human activities would have affected such changes.

In this research, I recollected the presence and absence data of 8 kinds of wild mammals in eastern China from 1573C.E.(Common Era) to 1949C.E. I chose this time period and region to avoid potential biases, because (*i*) eastern China was the main activity area of ancient Chinese people; (*ii*) during this period, China experienced a rapidly growing human population; (*iii*) a greater quantity of records of wild animals in regional gazetteers. The data were re-organized from scanned local gazetteers (county-level or prefectural-level) in the Chinese Digital Gazetteer Database (http://x.wenjinguan.com/) and *The Series of Rare Local Gazetteers in the National Library* (Fu *et al*. 2016). All 8 kinds of the mammals I studied are predators of distinctive body sizes, are widely distributed across gazetteer records, and are also easily distinguishable in traditional folk taxonomy. Furthermore, by performing new analytical techniques (see *Methods*), I reconstructed the spatial and temporal distribution patterns of 8 kinds of predators with both presence and absence records. The ultimate objective of this study was to quantitatively and specifically estimate how human socioeconomic activities have affected the distribution and extirpation of these 8 predators in eastern China.

## 2. Methods

### Overview

I chose the eastern part of China as the research area, and the temporal span 1573C.E.-1949C.E. as the research period, for there are rich amount of local gazetteers preserved, and the human population experienced a huge increase. I collected both the presence and absence data of each target mammal predator from local gazetteers and sorted them into 4 periods. In each period, I used the tool Inverse Distance Weighted (IDW) to reconstruct the distribution probability layers by ArcGIS 10.3, I named this procedure as “Interpolation of Presence & Absence Records in Same Periods”. Then I used 20km*20km grids to extract the continuous value in those layers and the human population density layers, to analyze the distribution changes, the relation between animals, and the influence of humans. (see *Fig. S1*)

### 2.1 Data collection

I collected the data including both presence records and absence records from the *Chinese Digital Gazetteer Database*(http://x.wenjinguan.com/) and *The Series of Rare Local Gazetteers in the National Library*(Fu *et al*. 2016). Previous studies used only presence points to reconstruct the presence and pseudo-absence(actually confused the real absence(0) with data missing(null) maps, then extracted the binary value in each grid to analyze with limited maths methods. Different from the above way, I used both presence and absence points to reconstruct the probability maps. Then I extracted the continuous value in each grid to analyze. Previous studies collect the presence records of the target animal, but in this research, I collected the valid animal diversity records, and acquired the presence and absence records of the target animal. For example, there are two animal records in different local gazetteers(see row 1 in *Fig.S1*), A recorded “Tiger, Leopard, Deer, Roe Deer …. Hedgehog, Vole, Ibex.” (Wolf not existed), and B recorded “Cattle, Horse … Wolf, Dog, Badger, Rabbit, Mouse”(Wolf existed). The previous studies would output “There were wolves in B” merely, but my study would output“There were wolves in B, and there were no wolves in A”(see row 2 in *Fig.S1*).

There are two kinds of invalid records I excluded. The first kind recorded only livestock, and the second kind did not record comprehensively but recorded only several large mammals roughly. To avoid the above invalid records, I filtered the gazetteer records by the following rule: a valid record should record any wild mammal of which the body weight is below 45kg(including those not analyzed in this research, like macaca monkeys, gibbons, or wild rabbit) (see *Table.S1*).

I select 8 kinds of wide-distributed predators, i.e. tigers, leopards, bears, wolves, foxes, civets, dholes, and mustelids, to research. And I match the folk animal kinds to the modern species in China(Smith & Xie 2008) (see *Table.S2*), the folk taxonomy animal kind can be matched to one or several modern scientific taxonomy species. I sorted tigers, leopards, and bears into the large/apex predator group, and the rest into the mesopredator group, according to the definition of megafauna, body mass≥45kg or occupying the relevant ecological niche (Roberts *et al*. 2001; Barnosky 2008; Boulanger & Lyman 2013; Villavicencio *et al*. 2016; Moleon *et al*. 2020).

### 2.2 Spatial processing

I selected 22 province-level regions in the eastern continent of China as the research area (excluding the Tibetan Plateau part in Sichuan and Yunnan) which roughly corresponds with the eastern region of the Heihe-Tengchong Line (see *Fig.1*), containing approximately 90% of the human population of China from ancient to modern eras (Hu 1935). Although the names and areas of many administrative divisions have changed, it is able to locate the data on modern maps by using the current local government locations to identify the corresponding locations of the local gazetteers.

**Fig.1.**
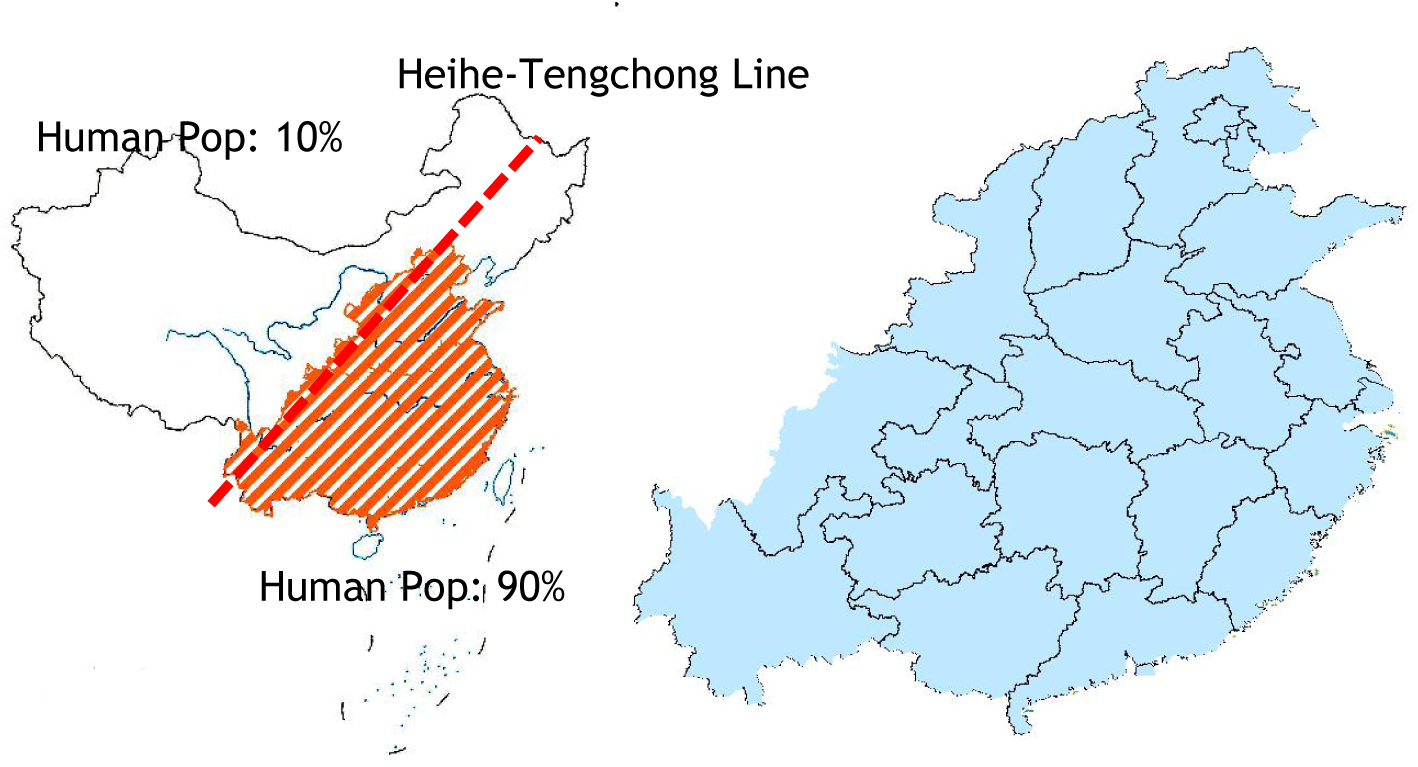
Research area (blue) in eastern China and the Heihe-Tengchong Line (red). I selected 22 province-level regions in eastern continent China as the research area (excluding Tibetan Plateau part in Sichuan and Yunnan) which roughly corresponds with the eastern region of the Heihe-Tengchong Line, containing approximately 90% of the human population of China from ancient to modern eras (Hu 1935).

**Fig.2.**
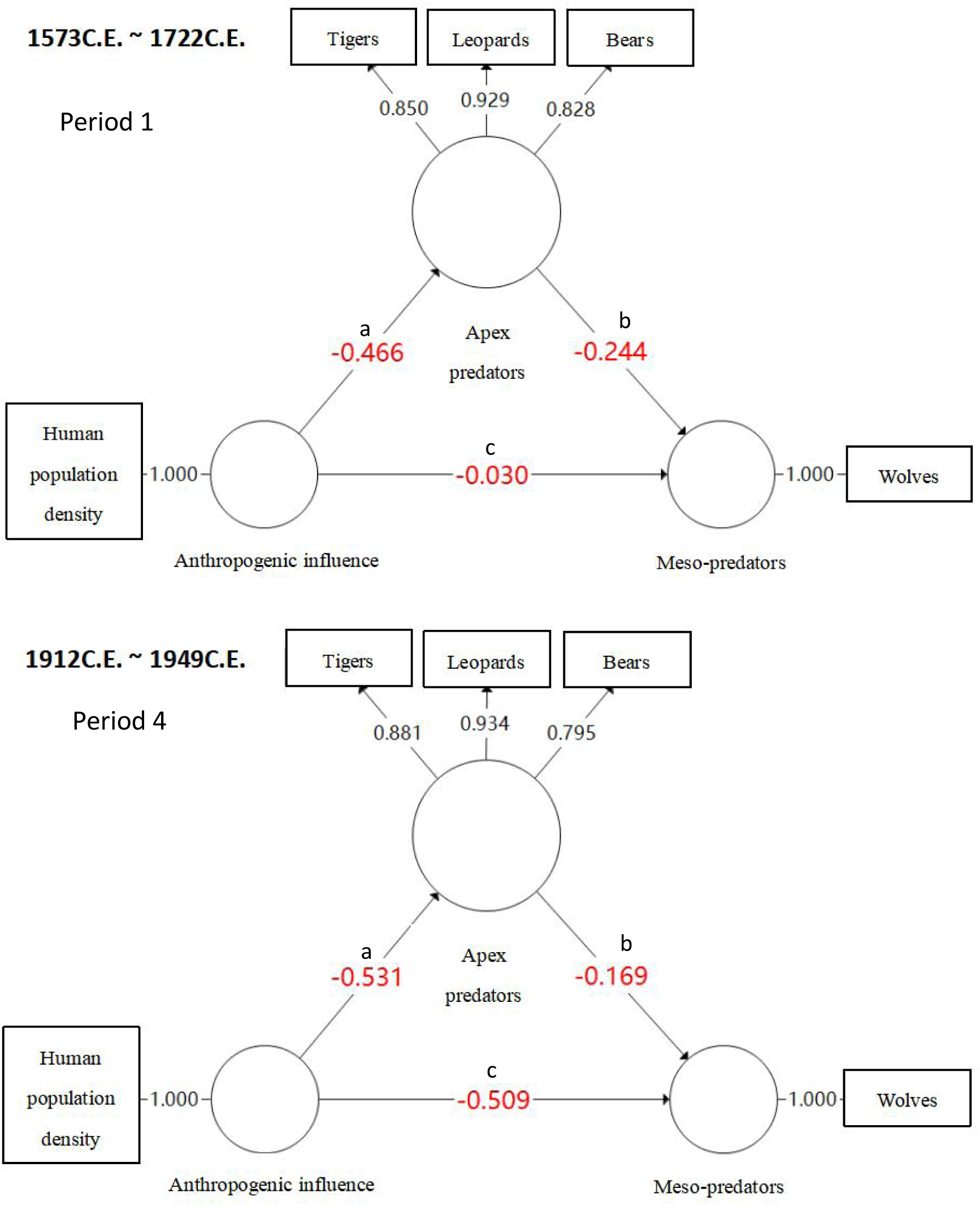
The PLS-SEM models. In the triangle, the arrow “a” is the Anthropogenic influence on Apex predators; the arrow “b” is the Apex predators’ influence on Meso-predators; the arrow “c” is the direct Anthropogenic influence on Meso-predators. The overall Anthropogenic influence on Meso-predators is: s=a*b+c. Exemplified by wolves in Period 1 and Period 4: (i)HPD exerted a slight negative direct influence: -0.030. HPD also exert a large positive indirect influence by kill the apex predators:(-.466)*(-0.244)≈0.114. The total HPD influence is 0.084, positive.HPD exerted a large negative direct influence:-0.509. (ii)HPD also exert a smaller positive indirect influence by kill the apex predators:(-.531)*(-0.169)≈0.090. The total HPD influence is -0.419, negative.

I used the Inverse Distance Weighted (IDW) tool to reconstruct the distribution based on the valid records using ArcGIS Desktop 10.3. This tool has been widely applied to species distribution reconstruction (Roberts *et al*. 2004; Hijmans 2012; Takahashi *et al*. 2014; Barbosa 2015; Areias-Guerreiro *et al*. 2016; Palacio *et al*. 2021). The interpolation value in each grid was generated by the nearest 12(default value) records, including presence records or absence records. The main assumption is that species are more likely to be found closer to presence points and farther from absences (i.e. spatial auto-correlation), and the local influence of points (weighted average) diminishes with distance (Palacio *et al*. 2021). The result value means the probability of animal occurrence, in [0,1]. Besides the accuracy of data, the smoothy probability value is continuous, so I can analyze the distributions much subtler.

### 2.3 Temporal processing

To keep the balance between the diachronical-spatial dimensions, I selected 1573-1949 as the research span, and sorted the valid points into 4 periods according to the reigning emperor, since many local gazetteers are labeled under the reigning emperor’s title without specifying the year, and adopted the later one when there is more than one record in the same location in one period. Period 1 covers the span of 1573C.E.-1722C.E., with 694 valid data points; Period 2 covers the span of 1723C.E.-1820C.E., with 706 valid data points; Period 3 covers the span of 1821C.E.-1911C.E., with 786 valid data points; and Period 4 covers the span of 1912C.E.-1949C.E., with 451 valid data points. These layers are cross-sections and based on both presence/absence points, so it is appropriate that spans are different in periods.

This research has excluded the modern distribution of mammalian predators because their distribution was greatly reduced as a consequence of a campaign by the Chinese government to hunt the “harmful beasts” since the 1950s.

There are few local gazetteers that were written before 1573C.E. and are still preserved today. That means the presence records (also absence records) are rare in this period. Besides, the Chinese human population experienced several fluctuations in history. It is inappropriate to infer the decline trend with the sample with these biases. So I did not analyze this period in this research.

### 2.4 Statistics

#### 2.4.1 Fishnet grids for overall surveys

I used the fishnet tool to build 10819 20km*20km grids (in WGS 1984 World Mercator projected coordinate system), to extract the data from the layers.

The significance test is not necessary for this research, because the statistics are not sampling surveys but overall surveys.

#### 2.4.2 Relations of predator distribution

I used Kendall’s tau-b coefficient to measure the relations between each predator in SPSS 22. This statistic is a measure of the strength and direction of associations between two Ordinal variables. Kendall’s tau-b treats the variables symmetrically and ranges in [-1,1], with the ± symbol reflecting the direction of correlation and the absolute value suggesting the strength of association.

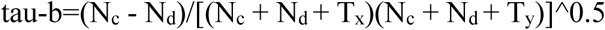

N_c_ is the number of concordant pairs, N_d_ is the number of discordant pairs. T_x_ & T_y_ are the number of tied pairs on each variable.

#### 2.4.3 Influence of anthropogenic factor on predator distribution

I added the human population density layers to make comparisons. I interpolated human population data for the last three periods based on historical demographic research. The population data in this study is referenced from Cao’s(2001) “History of the Chinese Population, Volume Five, Qing Dynasty” for historical population statistics from the Qing Dynasty to 1953C.E. The 1820C.E. data corresponds to the second period (1723C.E.-1820C.E.); the 1910C.E. data corresponds to the third period (1821C.E.-1911C.E.), and the 1953C.E. data corresponds to the fourth period (1912C.E.-1949C.E.) (Cao 2001). I imported the human population data into ArcGIS Desktop 10.3 and created a kernel density map using a 200km search radius. For the first period, where precise population statistics were unavailable, the study used the population density layer from the HYDE 3.2 database for the year 1720C.E. as a substitute (Goldewijk et al. 2017).

I measured the influence of human population density(HPD) on predator distribution between each predator by Somers’ d_yx_ statistic in the software SPSS 22. This statistic is a measure of the influence strength of an ordinal variable on the other ordinal variable. The d_yx_ ranges in [-1,1], with the ±symbol reflecting the direction of influence and the absolute value reflecting the strength (Somers 1962).

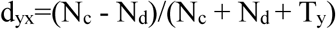

N_c_ is the number of concordant pairs, N_d_ is the number of discordant pairs. T_y_ is the number of tied pairs on the dependent variable.

#### 2.4.4 Terrain factors

This research incorporated the elevation layer and slope layer (both are from Loess Plateau SubCenter, National Earth System Science Data Center, National Science & Technology Infrastructure of China (http://loess.geodata.cn)). The entire grids were approximately divided into five groups based on elevation (in meters): alt1([-5, 58], n=2169 grids), alt2((58, 249], n=2157 grids), alt3((249, 670], n=2162 grids), alt4((670, 1241], n=2166 grids), and alt5((1241, 4414], n=2165 grids). Subsequently, I used charts to display the variations in different animal species and human population within each elevation range. The structural equation models analyzing the complex causal relationships among large/apex carnivores, medium-sized carnivores, and human population density.

I used PLS-SEM to analyze the complex causal relationships among the human population, large carnivores, and medium-sized carnivores. In the first step, I measured the latent variable of the overall large carnivores(reflecting the completeness of the food chain) using the three kinds of large carnivores as reflective indicators. In the second step, I conducted statistical analyses with human population density as the independent variable, the overall large carnivores as the mediating variable, and each kind of medium-sized carnivores as the dependent variable. Because the large carnivores have predatory and competitive relationships with the medium-sized carnivores in the food chain, and humans hunted both large and medium-sized carnivores, I analyzed whether the large carnivores mediated the impact of population density on medium-sized carnivores and the resulted masking effect.

I chose PLS-SEM instead of the traditional CB-SEM, for the latter requires a normal distribution of data. Compared to the traditional covariance-based structural equation modeling (CB-SEM), partial least squares structural equation modeling (PLS-SEM) does not require the data to follow a normal distribution as it employs a non-parametric estimation method. In this study, the IDW data did not exhibit a normal distribution, and there was a large amount of grids with values of 0 or 1 (representing extreme cases of 0% or 100% probability distribution). These grids made it unsuitable to transform the data into a normal distribution through standardization (which requires a larger sample size in the middle range and fewer extreme values at both ends).

## 3. Result

### 3.1 The change of distribution area

I reconstructed the distribution of **target predators and human population density** (HPD) (see *Table.1*). In the large/apex predator group, the distribution of the two largest predators, tigers and bears retreated, while the smaller one, leopard, remained almost unchanged; by contrast, the distribution of predators in the mesopredator group expanded in varying degrees (see *Fig.3 & Table.S3*).

**Fig.3.**
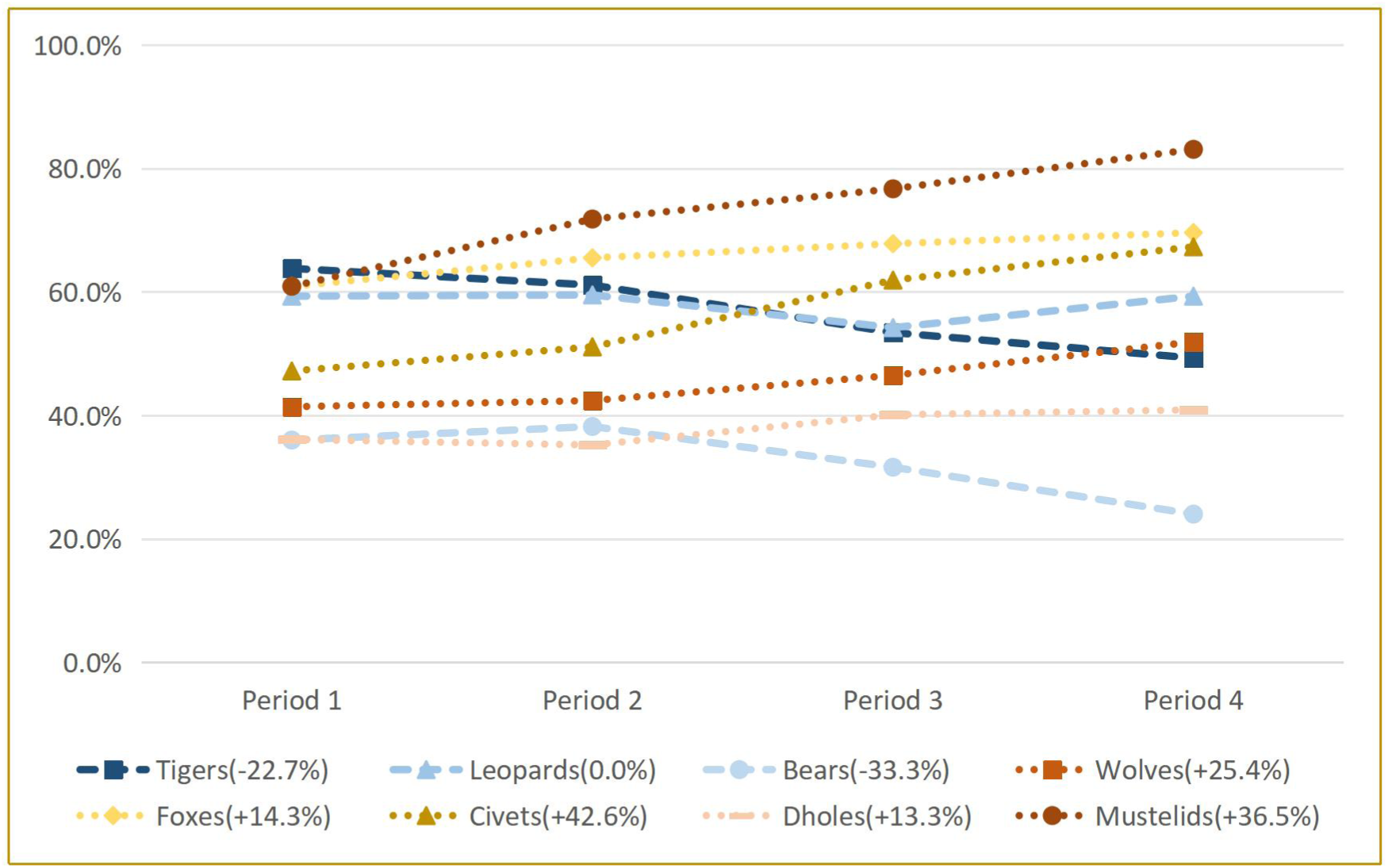
The change of average probability of predators from Period 1 to Period 4, after the carnivores’ name are the total change proportion from Period 1 to Period 4: (S_Period 4_-S_Period 1_)/S_Period 1_.

**Table.1.**
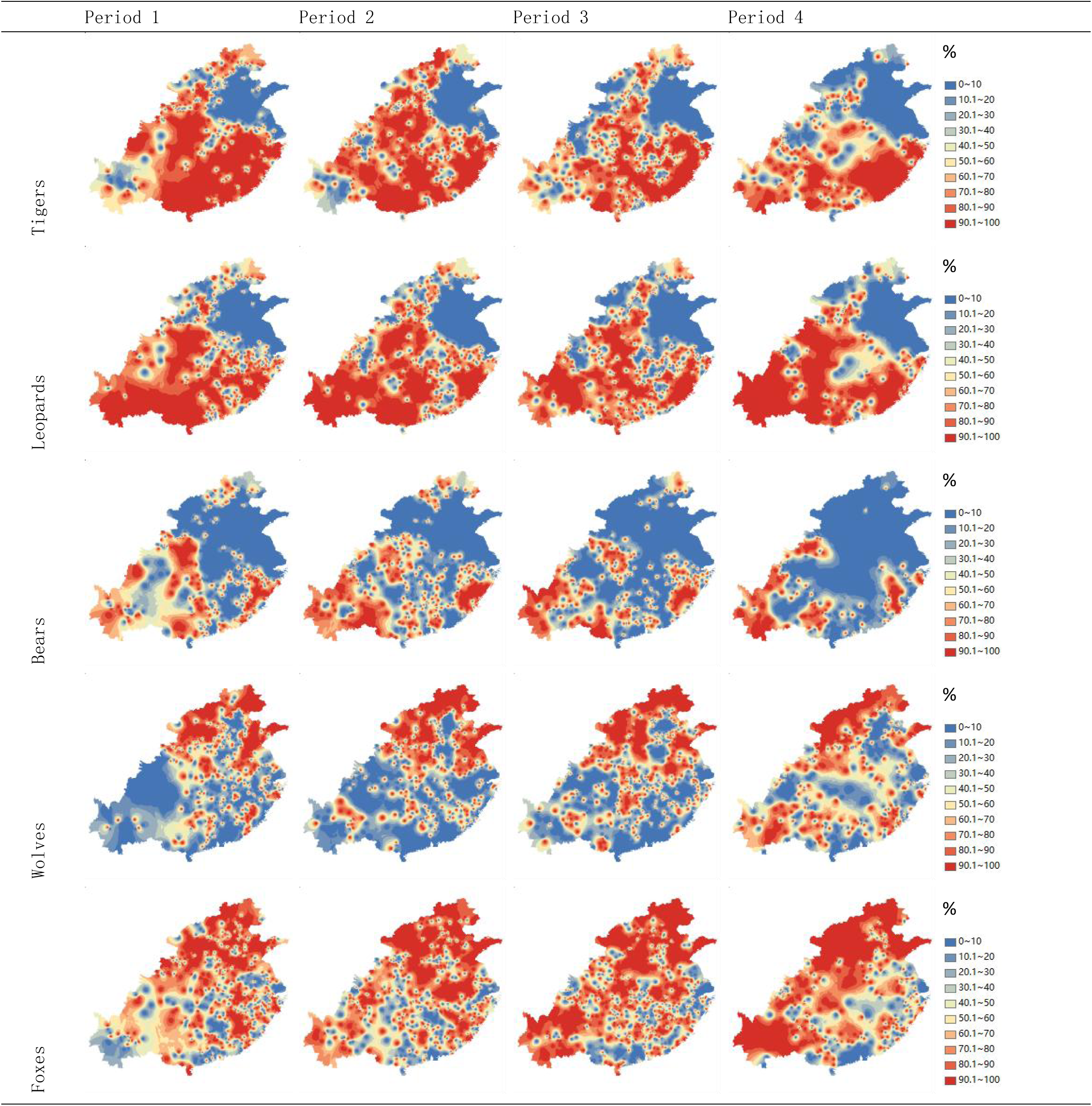
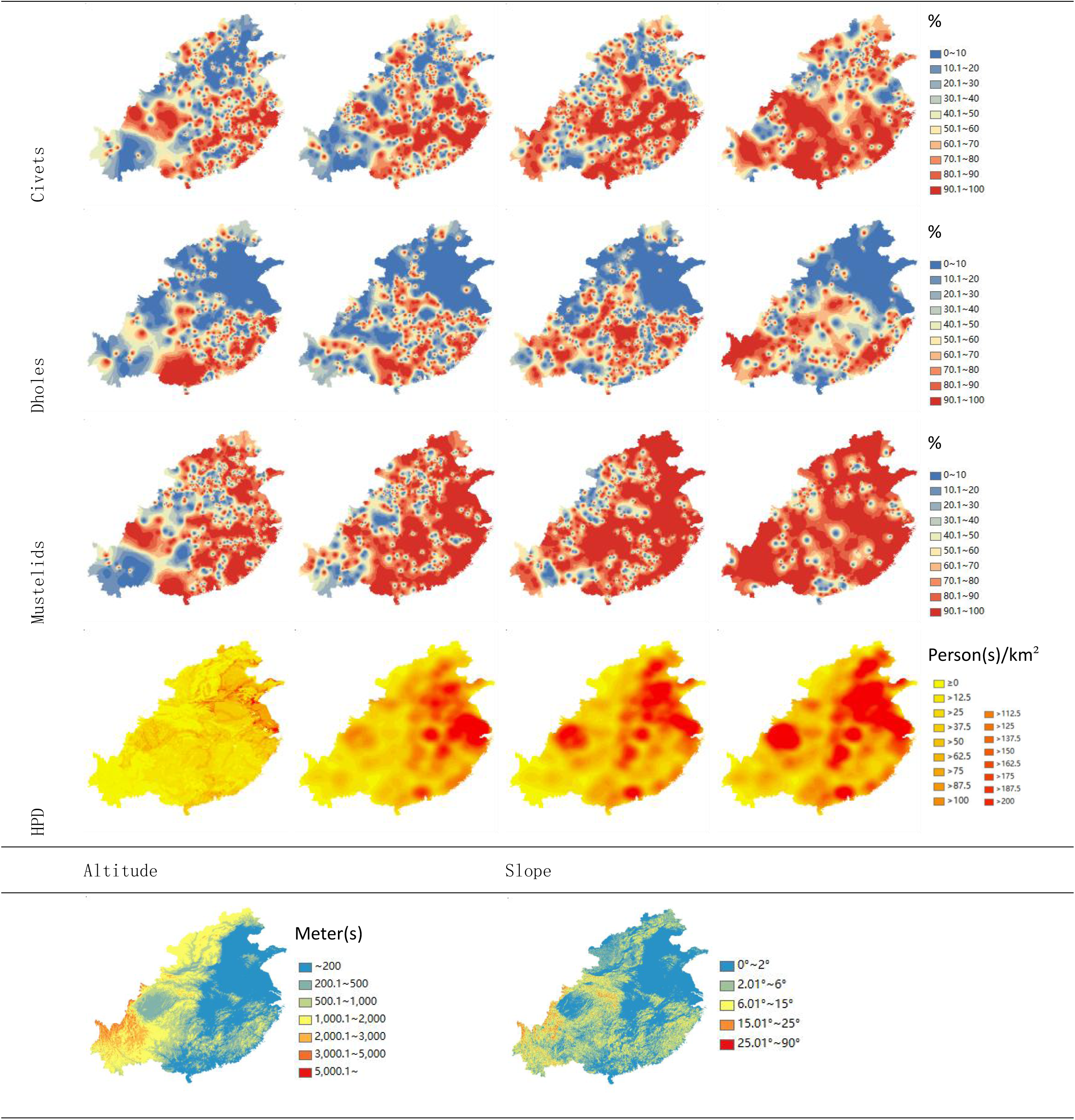
The distribution of carnivores, human population density(HPD), and Altitude/Slope.

Based on the analyses of elevation and slope, the average distribution of various carnivorous mammals overall shows an increasing trend in both elevation and slope (see *Fig.4*). Among them, the four periods of the large carnivores and dholes are all higher than the overall average elevation and slope of the entire research area, and they exhibit an increasing trend. In contrast, the average elevation and slope of the other medium-sized carnivores were mostly lower than the overall average values initially but increased to above the average later.

**Fig.4.**
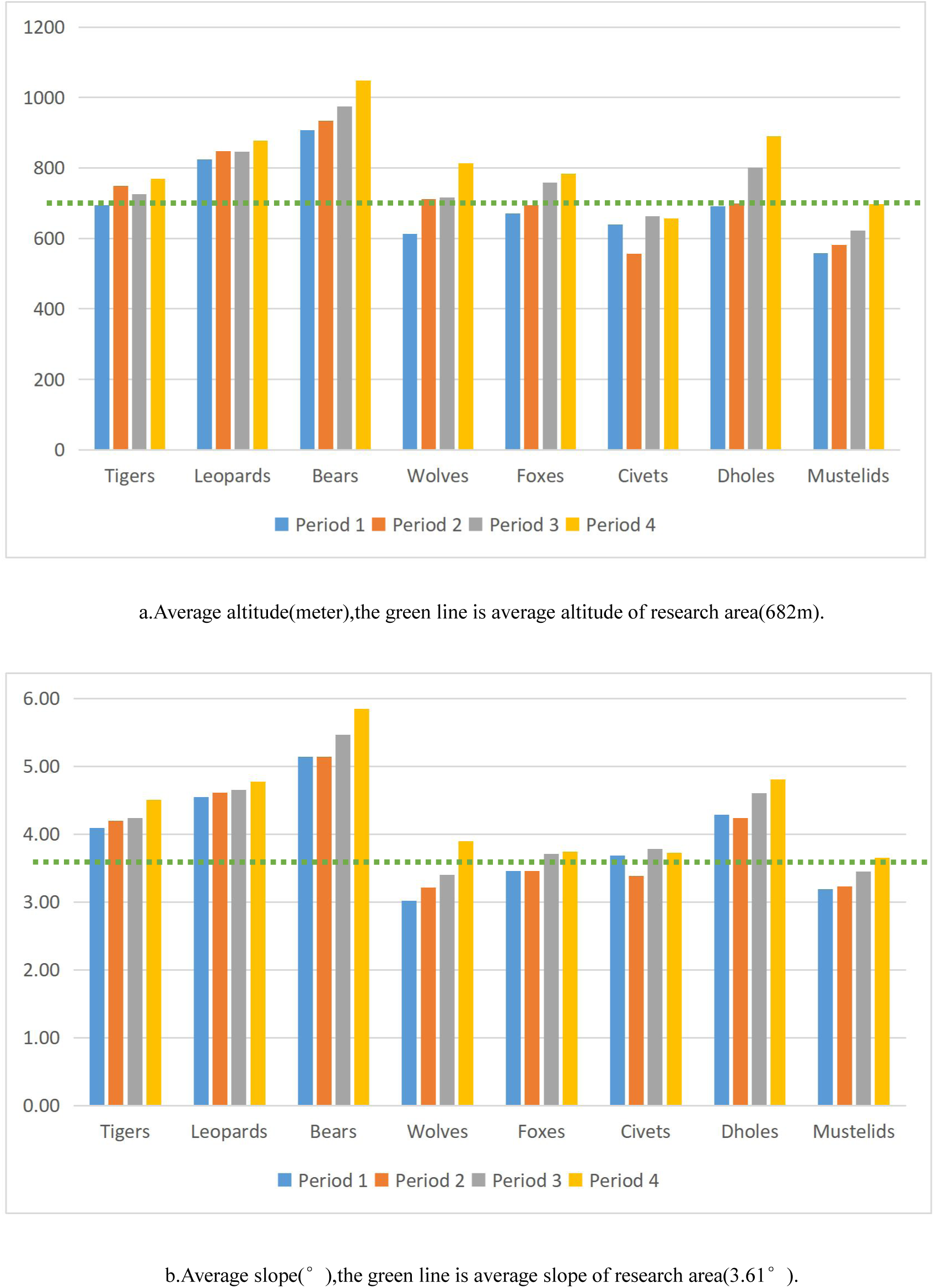
The average distribution altitude and average distribution slope of each kind of carnivores in each period.

The grouped data on altitude suggests that: 1) generally, in lower altitude regions, the distribution of large carnivores decreased obviously, while the distributions of medium-sized carnivores increased obviously(civets), kept almost stable(foxes, dholes, and mustelids), or decreased obviously(wolves); 2) generally, in higher altitude regions, the distributions of large carnivores kept almost stable, while the distributions of medium-sized carnivores increased obviously(see *Fig.S2, Fig.5, Table.S4)*.

**Fig.5.**
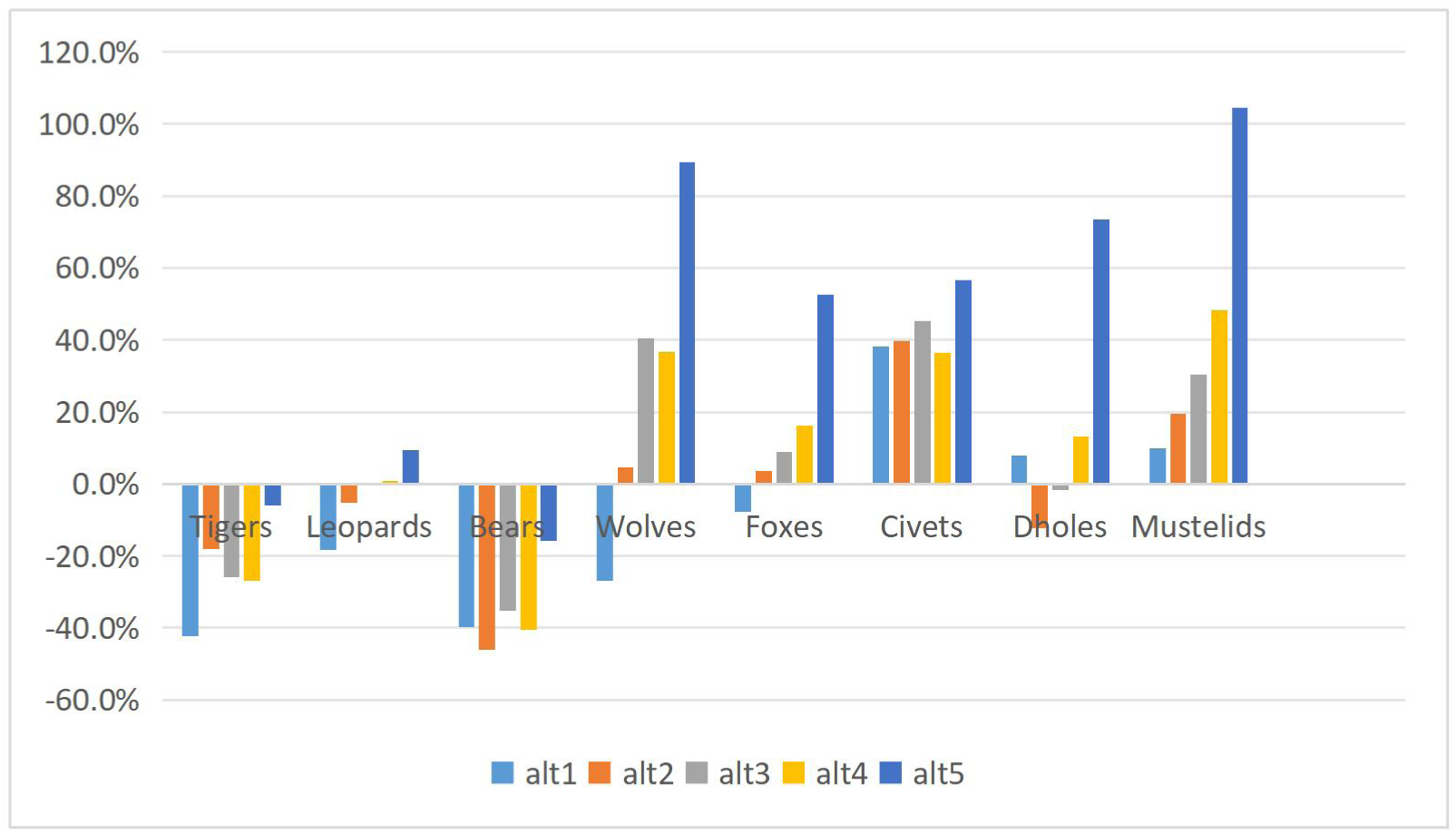
The carnivore distribution change proportions from Period 1 to Period 4 in each altitude group, i.e.: (S_Period 4_-S_Period 1_)/S_Period 1_.

### 3.2 Relations between each predator and others

According to the relations of distributions between predators (see *Fig.S3 & Table. S5*), the predators in the large/apex predator group displayed strongly positive correlations with each other in all periods**, suggesting that these species have similar environmental requirements**. In the mesopredator group, wolves and foxes were highly correlated, and the correlations between wolves/foxes and large/apex predators were negative overall; while for dholes and civets, they were positively correlated with the large/apex predators and each other, but weakly negative with wolves/foxes overall, suggest that they were adapted to coexisting with large/apex predators; for mustelids, the values with leopards and bears were negative in earlier periods but turn positive later, and the value with other predators(including tigers) were positive.

### 3.3 Anthropogenic influence on predator distributions

The results (see *Fig.6 & Table. S6*) show that HPD exerted a negative influence upon the distributions of each large predator during each period, the influences on tigers, leopards, and bears kept are clearly negative throughout. But for mesopredators: (1)wolves, foxes, and civets were slightly positive in early periods, but became negative in later periods; (2)dholes were negative in all periods, like large predators; and (3) mustelids were strongly positive in period 1 to period 3, but turn slig<u>htly</u> <u>negative</u> in period 4.

**Fig.6.**
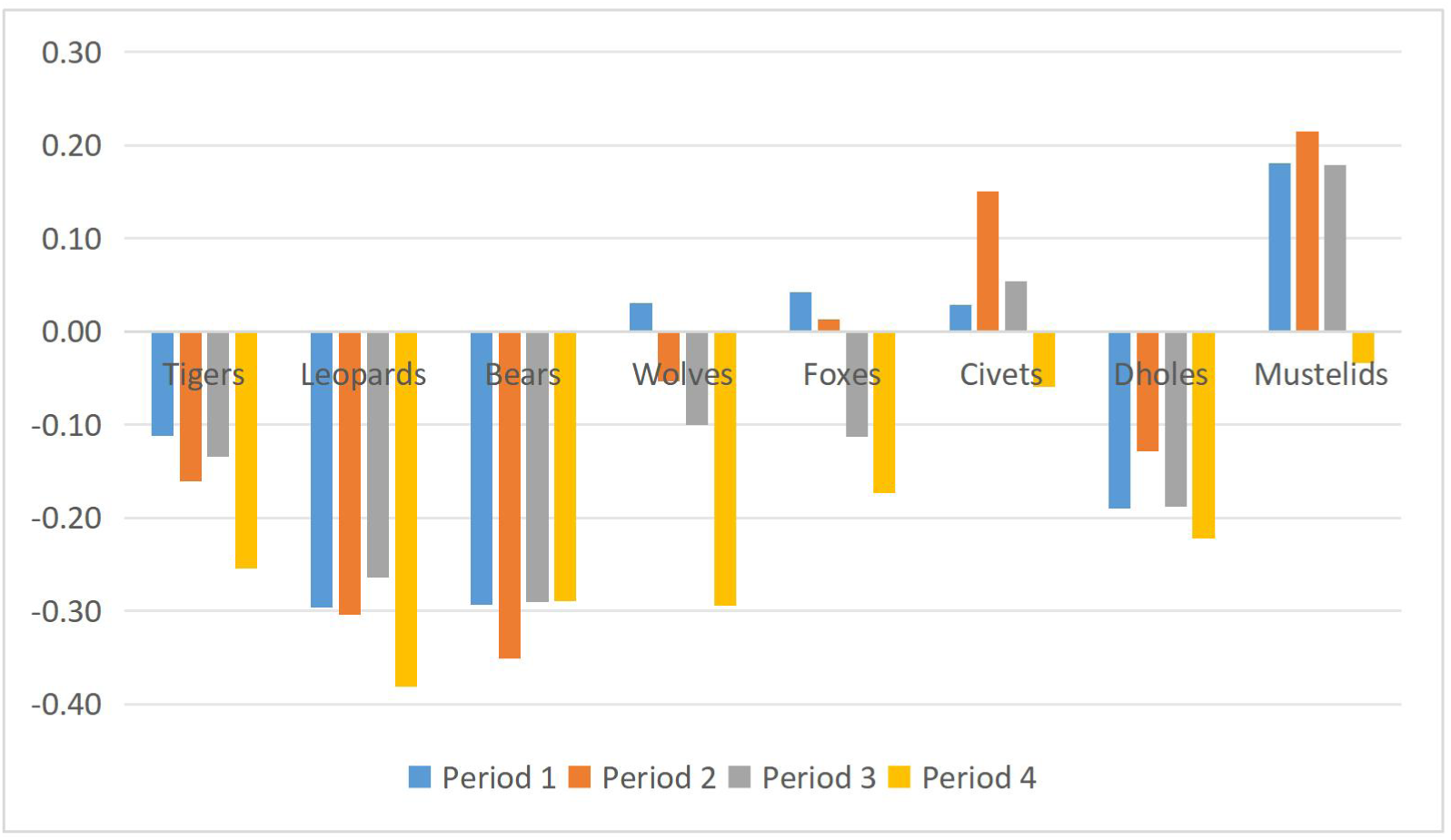
HPD Influence on predator distribution (Somers’ dyx)

### 3.4 Complex causual relationships in PLS-SEM

According to the analysis(see *Table.2*), in the triangular relationship of “anthropogenic influence(human population density) - the overall Apex predators(large carnivores) - each of Meso-predators(medium-sized carnivores)” across four periods, human population density has consistently exerted a strong negative impact on the overall large carnivores (a).

**Table.2.** The results of PLS-SEM.

|  |  | a.HPD→Apex | b.Apex→Meso | c.HPD→Meso<br>(directly) | s.HPD→Meso<br>(total) |
| --- | --- | --- | --- | --- | --- |
| Wolves | Period 1 | -0.466 | -0.244 | -0.030 | 0.084 |
|  | Period 2 | -0.477 | -0.298 | -0.218 | -0.076 |
|  | Period 3 | -0.481 | -0.211 | -0.213 | -0.112 |
|  | Period 4 | -0.531 | -0.169 | -0.509 | -0.419 |
| Foxes | Period 1 | -0.466 | -0.143 | -0.067 | 0.000 |
|  | Period 2 | -0.476 | -0.232 | -0.151 | -0.040 |
|  | Period 3 | -0.481 | -0.107 | -0.208 | -0.156 |
|  | Period 4 | -0.529 | -0.278 | -0.348 | -0.201 |
| Civets | Period 1 | -0.460 | 0.390 | 0.155 | -0.025 |
|  | Period 2 | -0.472 | 0.250 | 0.256 | 0.138 |
|  | Period 3 | -0.476 | 0.361 | 0.173 | 0.002 |
|  | Period 4 | -0.529 | 0.326 | 0.065 | -0.108 |
| Dholes | Period 1 | -0.460 | 0.678 | 0.032 | -0.280 |
|  | Period 2 | -0.467 | 0.686 | 0.140 | -0.181 |
|  | Period 3 | -0.474 | 0.648 | -0.001 | -0.309 |
|  | Period 4 | -0.529 | 0.683 | 0.032 | -0.329 |
| Mustelids | Period 1 | -0.467 | 0.032 | 0.225 | 0.210 |
|  | Period 2 | -0.478 | -0.039 | 0.223 | 0.238 |
|  | Period 3 | -0.479 | 0.232 | 0.340 | 0.229 |
|  | Period 4 | -0.530 | 0.075 | -0.032 | -0.071 |

For wolves and foxes, the direct impact of human population density (c) was consistently negative, and it became stronger (i.e. the absolute values increased) as the HPD increased in different periods. The impact of large carnivores was also consistently negative (b). In summary, the overall impact of HPD (s) initially was positive or nearly neutral, but later turned negative and became increasingly stronger.

For civets and dholes, the direct impact of HPD (c) was consistently weakly positive or nearly neutral, and the impact of large carnivores was strongly positive(b). In summary, the overall impact of HPD (s) fluctuates between weakly positive and weakly negative effects for civets, while it has consistently been negative for dholes.

For mustelids, the direct impact from HPD (c) was positive in the first three periods but became weakly negative in the fourth period. The impact of large carnivores (b) was weakly positive in the first period, weakly negative in the second period, and strongly positive in the third and fourth periods. The overall impact of HPD (s) was initially strongly positive but turned weakly negative in the fourth stage.

## 4. Discussion

### 4.1 Discussion on procedure

In my procedure “Interpolation of Presence & Absence Records in Same Periods”, the reconstruction of distributions using both presence and absence data is more accurate than previous studies (Zhou & Zhang 2013; Li *et al*. 2015; Turvey *et al*. 2015; Wan *et al*. 2019; Teng *et al*. 2020) with have only used presence data (see *Fig.7*), especially with the last observation procedure (see *Table.S7 & Table.3*). It overcomes several disadvantages of previous studies:

1. Sampling bias: The distribution of local gazetteers is not uniform in either space or time. If only considering the presence (1) records, it will greatly confuse the absence (0) areas and unknown(null) areas. There are few gazetteers in the area/period with social turbulence or few human population, causing the presence records to be few there/then (*Fig.7.a,*). I fixed this bias and interpolated the unknown areas by the IDW.
2. New data: if there are new data, the reconstruction might change obviously (e.g. there are many more newly found records in Wen(2018)’s new research than his previous research (Wen 2009) then the reconstruction changed obviously), for the last observation precedure, it would lead to inverse interpolation. While in my procedure, new data will subtlize the reconstruction.
3. Subjective: how large an area (circle or square, 50km*50km or 100km*100km, etc.) a data point represents is determined by the subjectivity of different researchers (Zhou & Zhang 2013; Turvey *et al*. 2015; Wan *et al*. 2019; Teng *et al*. 2020), leading to different result of distribution area, but in this research, it is replaced by objective interpolation of IDW.
4. Monotonic recession prejudged: Last observation precedure can not reflect the expansion or recovery of animal distribution, but only monotonic recession, it is a kind of prejudge. In contrast, the expansion or recovery can also be noted in my procedure.

**Fig.7.**
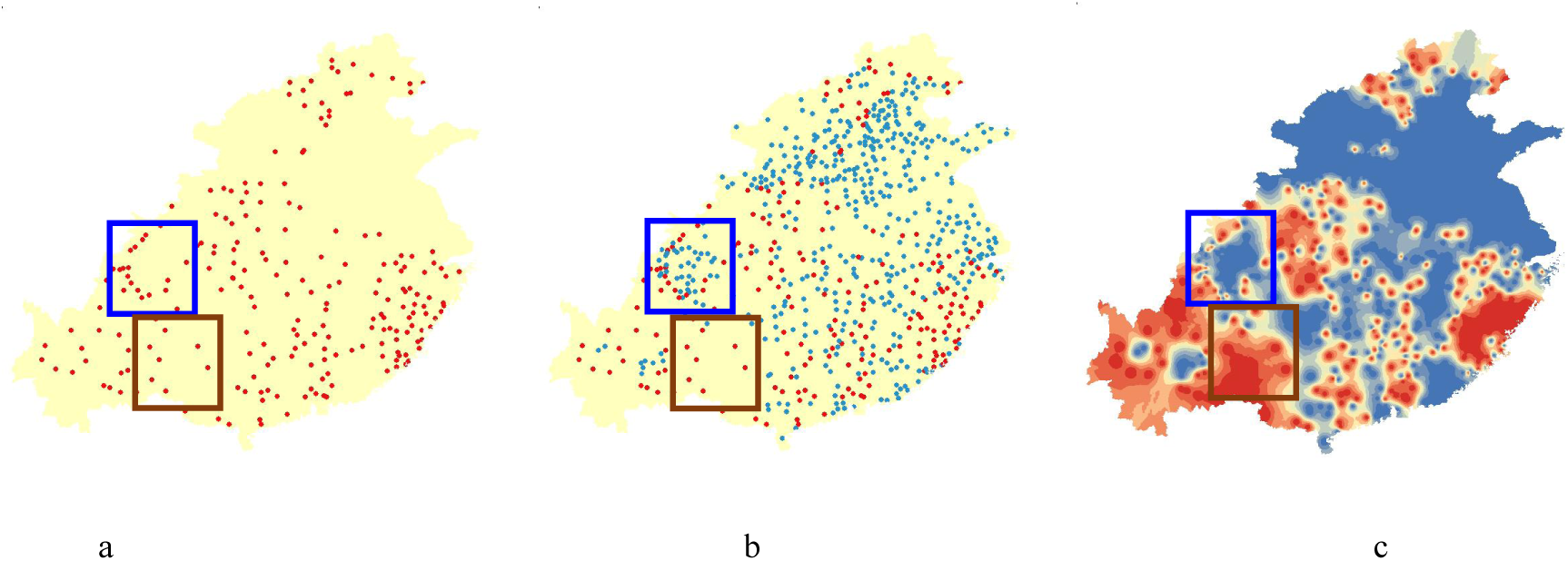
Criticizing the presence-only method a.The bear presence records(red points) in Period 2, both frames are almost empty; b. The bear presence records(red points) and absence records(blue points) in Period 2, there are many absence records in the blue frame, but few in the brown frame; c. The IDW reconstruction of bear distribution based on b., high-probability area(red) and low-probability area(blue), the blue frame is fulfilled by low-probability area but the brown frame is fulfilled by high-probability area.

Above all, there are several biases in previous studies, and the procedure in this study mended them.

I exemplify the contrast of the two procedures by the distribution of wolves(see *Fig.S2 & Table.3*). The results in the last observation precedure show a retracting trend, but the result values are much different between 100km*100km square grids and 50km*50km square grids. In contrast, the result in the IDW shows an expanding trend.

**Table.3.** Different results of IDW, Last Observation(100km*100km) & Last Observation(50km*50km)

|  | IDW | Last Observation<br>100km*100km | Last Observation<br>50km*50km |
| --- | --- | --- | --- |
| Period 1 | 41.4% | 73.6% | 31.8% |
| Period 2 | 42.4% | 67.1% | 28.3% |
| Period 3 | 46.5% | 61.9% | 24.2% |
| Period 4 | 51.9% | 34.4% | 11.3% |

### 4.2 Discussion on results

#### 4.2.1 Distribution of carnivores

The distribution of wild animals is significantly shaped by anthropogenic disturbances throughout the long history of China (Chatterjee *et al*. 2012).

The distribution of large carnivores decreased or kept almost stable. I found that the distribution of the two largest carnivores, i.e. tigers and bears, retreated over successive periods, supporting similar views of previous studies (He 1993; 1997; Wen 2009; 2013; 2018; Wan *et al*. 2019; Teng *et al*. 2020), and reflecting the recession of the eco-system over the same time frame. Leopards are the smallest one in large carnivores and also contain the larger medium-sized carnivores: cloud leopards. The distribution of leopards kept almost stable, within the area in the lowest altitude group(alt1) decreased slightly but the area in the highest altitude group(alt5) increased slightly.

The distribution altitudes/slopes of large carnivores were consistently higher than the average in the research area and were increasing, suggesting a retraction towards higher altitudes (mountainous areas). Fisher and Shanee et al. have pointed out that for many animals currently concentrated in mountainous regions, the lower altitude areas are their most suitable habitats, however, human activities in these lower altitude areas compressed these animals’ living spaces to the higher altitude, suboptimal habitats on the edges of their original suitable habitats. This study supports this viewpoint (Fisher 2011, Shanee et al. 2014).

The decline trend of bears was severe than that of tigers. but the bears are still living in several protected areas in the research area, while tigers are already extirpated. I infer that the reason is the different home range sizes. Tigers, with large home ranges, would appear in wide sink-habitat out of source-habitat and be exposed to anthropogenic dangers; while bears’ habitats were majorly source-habitat, and they would survive when the small range habitats are preserved.

Not to be noted by previous studies, I find the distributions of mesopredator expanded in eastern China, matching the mesopredator release phenomenon, which suggests that the population of mesopredator would grow rapidly after the constraints from large/apex predators were withdrawn(Prugh *et al*. 2009). The distribution altitude/slope of medium-sized carnivores shifted from being lower than the average to approaching or higher than the average, suggesting an expansion of their distribution from lower altitude/slope areas to higher altitude/slope areas.

The results of mesopredators are not biased by data collection, because the distribution of other medium-sized mammals(2 kinds of primates and 3 kinds of medium-sized deer) decreased (Hu 2022).

I also provide a potential expression of the misunderstanding that wolves are not distributed in southern China. In numerous previous studies, wolves were once believed to be distributed only in the northern regions and the Qinghai-Tibet Plateau of China. However, Wang et al. clarified this misunderstanding by collating domestic specimen records and ecological research, confirming that wolves were also once widely distributed in various southern provinces of China (Wang et al. 2016). The evidence from this study supports Wang et al.’s viewpoint. It also indicates that the distribution of wolves in southern China undergoes a process of expansion, I speculate that it caused difficulties for early Western scholars to collect wolf specimens in southern China, thereby causing the misunderstanding.

#### 4.2.2 The relationship among carnivores

##### 4.2.2.1 Mesopredator release

The results match the changes of habitat overlaps in the process of mesopredator release. Newsome et al (2017) proposed the existence of three zones/stages during the process of mesopredator release: Zone A, the area monopolized by the large predators, where large predators are present but mesopredators are absent; Zone B, the area shared by large predators and mesopredators, where both present; Zone C, the area monopolized by mesopredators, where large predators have disappeared but mesopredators are still present. As HPD increased, the grouped altitude charts show the transition from Zone A to Zone B and from Zone B to Zone C. This study also identified a fourth zone/stage, Zone D, where mesopredators disappear as well when human population density was very high, where neither large nor mesopredators are present.

Exemplified by tigers and wolves, in the highest altitude group(alt5), the distribution of tigers just decreased slightly while the distribution of wolves increased obviously, corresponding to the transition from Zone A to Zone B. In the middle three groups, the distribution of tigers decreased and the distribution of wolves increased, corresponding to the transition from Zone B to Zone C. In the lowest altitude group(alt1), as the population becomes highly dense, both tigers and wolves decreased, corresponding to the transition from Zone C to Zone D.

It can explain why the relationships between large carnivores and wolves/ foxes/mustelids turned from negative to positive: the new co-exist area (Zone B expanded and replaced Zone A) and the new co-non-exist area (Zone D expanded and replaced Zone C) were larger than the new area where only large carnivores decreased (Zone C expanded and replaced Zone B).

I provide a new case about the switchable role of wolves. The body mass of wolves is transitive between apex predator and mesopredator. Although wolves are attributed to the role of a retreating apex predator in many studies (Prugh *et al*. 2009; Newsome *et al*. 2017), my results are more consistent with research conducted in Sikhote, Russia (Miquelle *et al*. 2005) where wolves are viewed as mesopredators being constrained by tigers. This difference shows that the classification between apex and mesopredator depends not only on species (Wallach *et al*. 2015), but also on the local eco-system.

##### 4.2.2.2 Predator guild

I confirm the predator guild among tigers, leopards, and dholes existed in long temporal span, and find out that bears (as a kind of large carnivore) and civets (as a kind of medium-sized carnivore) are probably also in the guild. Although dholes are a kind of mesopredator with a middling body size and its high-probability area expanded slightly, the distribution was strongly positively correlated with the large/apex predators, and only slightly positive with the other mesopredators. As a kind of mesopredator, dholes would be supposed to suffer from the constraints of apex predators, but recent studies in South Asia and Southeast Asia revealed that dholes can in general peacefully co-exist with tigers and leopards (Johnsingh 1992; Karanth & Sunquist 2000; Wang & Macdonald 2009; Jenks *et al*. 2012; Hayward 2014; Rayan & Linkie 2016), and thus form a relationship as a “predator guild”. This study supports the view of such a predator guild and supports it over a longer temporal span than previous studies. The relationship between dholes and bears is also strongly positively correlated, but I did not find any relevant predator guild study. So I infer that the reason is that the environmental demands of bears are similar to those of tigers and leopards, as the cause of such similar co-existing relations. Besides, still not noticed by previous research, civets were similar to dholes, but the positive relations with large carnivores were weaker than those of dholes.

#### 4.2.3 Influence of HPD

I quantified the specific negative impact of HPD on large carnivores, demonstrating the decline in the distribution of large carnivores under human interference. For medium-sized carnivores, the negative impact of HPD increased. The changes indicate a trade-off effect: in regions with moderate HPD, the positive impact of expelling large carnivores compensated for the direct negative impact o<u>f</u> humans’ killing medium-sized carnivores.

The PLS-SEM analyzed the relation between human population expansion and the mesopredator release. Wolves and foxes have been subject to sustained negative impacts from large predators, and the disappearance of large predators has reduced the survival pressure on wolves and foxes. Their distribution expansions were a typical instance of the mesopredator release effect. The masking effect also confirms the trade-off effect mentioned above. The trade-off effects of dholes, civets, and mustelids were not typical, as they are influenced positively or fluctuated from positively to negatively by large carnivores and HPD. Besides the predator guild effect in dholes and civets, I speculate that these three kinds of animals also benefit directly from human impact, such as providing more food sources like poultry and rodents.

In contemporary ecological research, red foxes (foxes in this research), wildcats (civets in this research), and stone martens (mustelids in this research) are common urban and suburban animals in Europe, and they are adapted to the human environment (Recio et al. 2015). It is similar to the results of this study on civets and mustelids. Dholes are not found in Europe, and the different situation of foxes in this study may be due to the prohibition of fur hunting in the modern.

## 5. Conclusion and Prospect

My study aimed to correct the existing procedures for reconstructions of historical distribution based on local gazetteers by adopting the use of both presence and absence data. Based on my reconstructions, I provide the direct proofs of human disturbance on mammal distribution in China, and extend support for the mesopredator release hypothesis and the predator guild hypothesis.

I have also collected records for other wild animals from local gazetteers in China. Based on the collection, I have built the Database of Wild Mammal Records in Chinese Local Gazetteers and are building the Database of Wild Bird Records in Chinese Local Gazetteers. I aim to continue relevant studies using these databases in the future.

## Acknowledgements

Ni, Ming provided advice to improve the figures and the tables. Hao, Chunhui provided advice to improve the introduction. Liu, Hanlun introduced the concept “mediating effect” to the author Hu, Kaijin.

## Supplementary Information

**Fig.S1.**
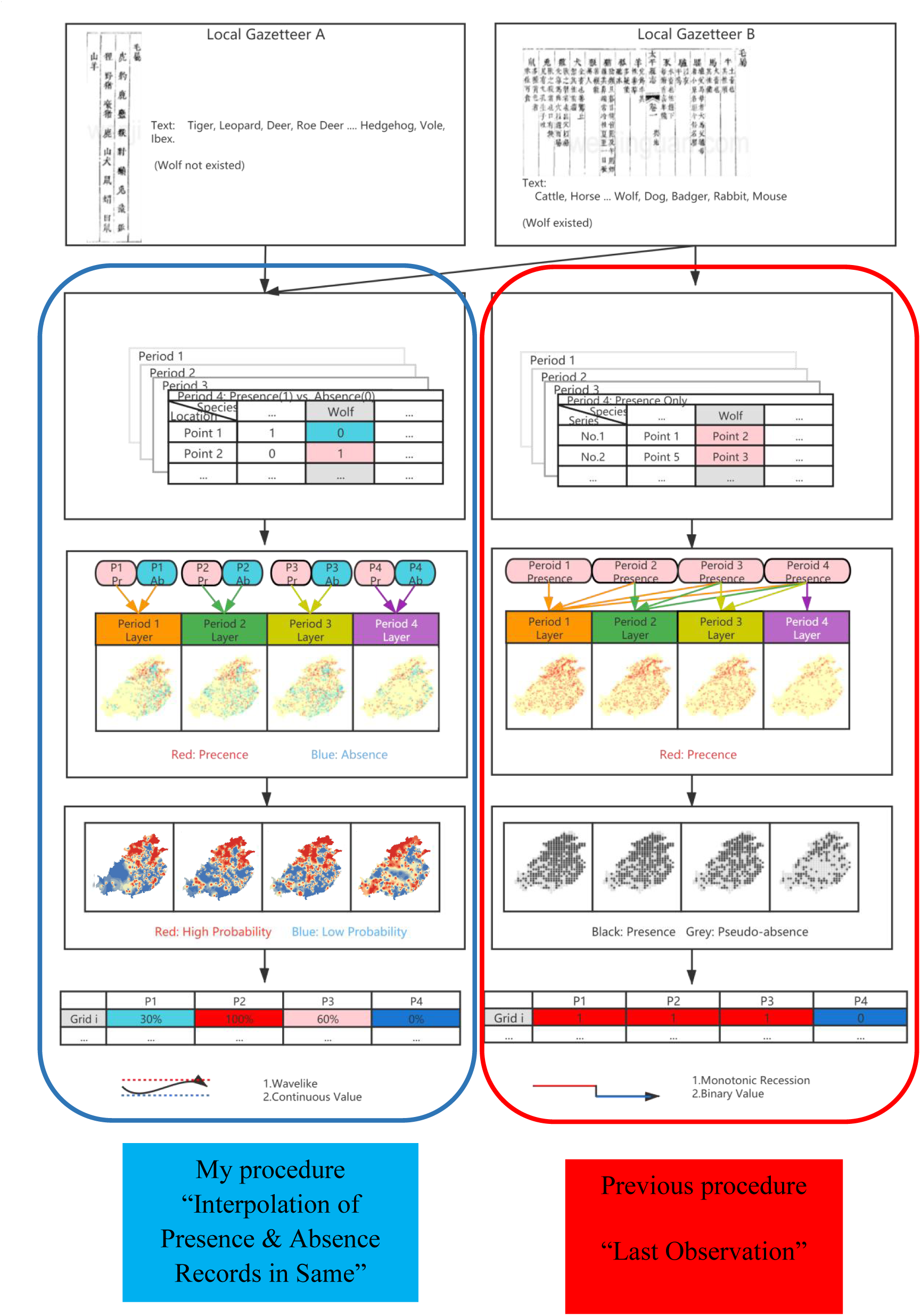
Flow Chart of the procedure “Interpolation of Presence & Absence Records in Same”.

**Fig.S2.**
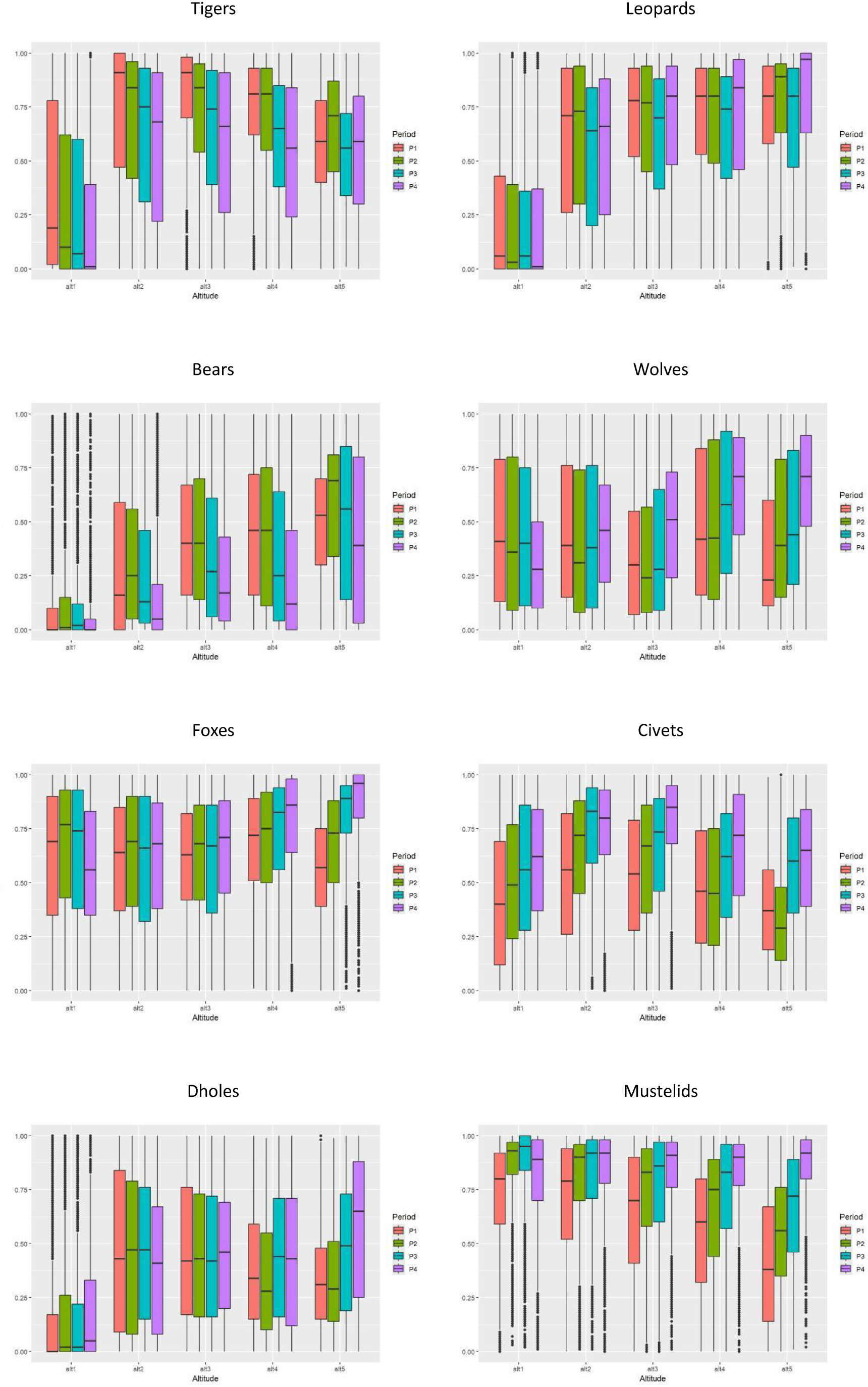
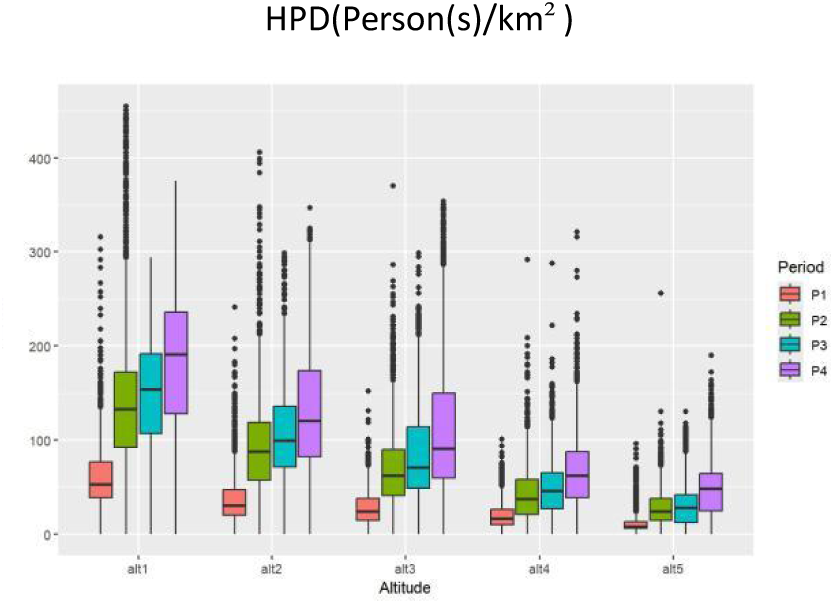
The carnivore distribution in each altitude group in each period.

**Fig.S3.**
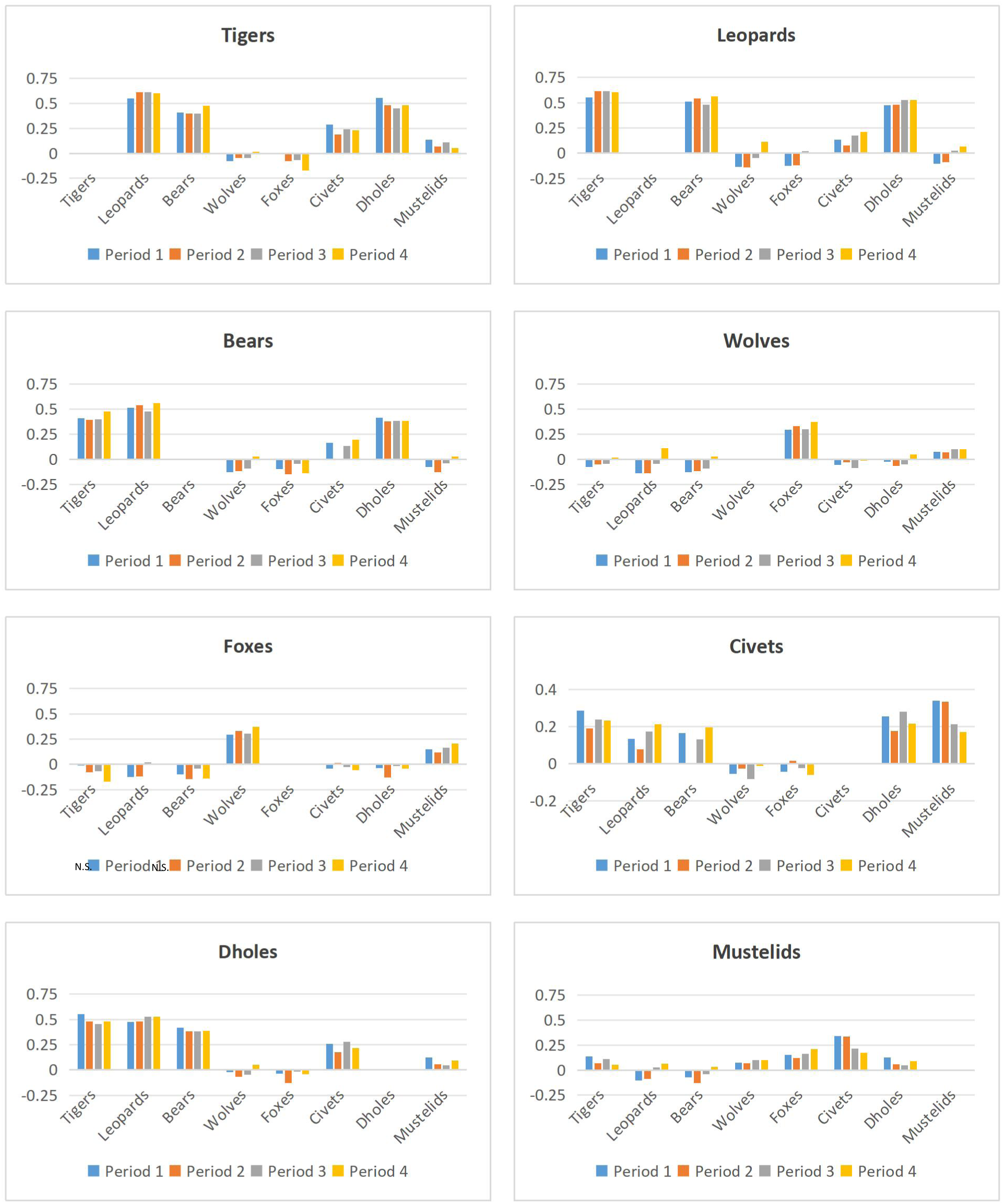
Relations between each predator and others (Kendall’s tau-b)

**Table.S1.**
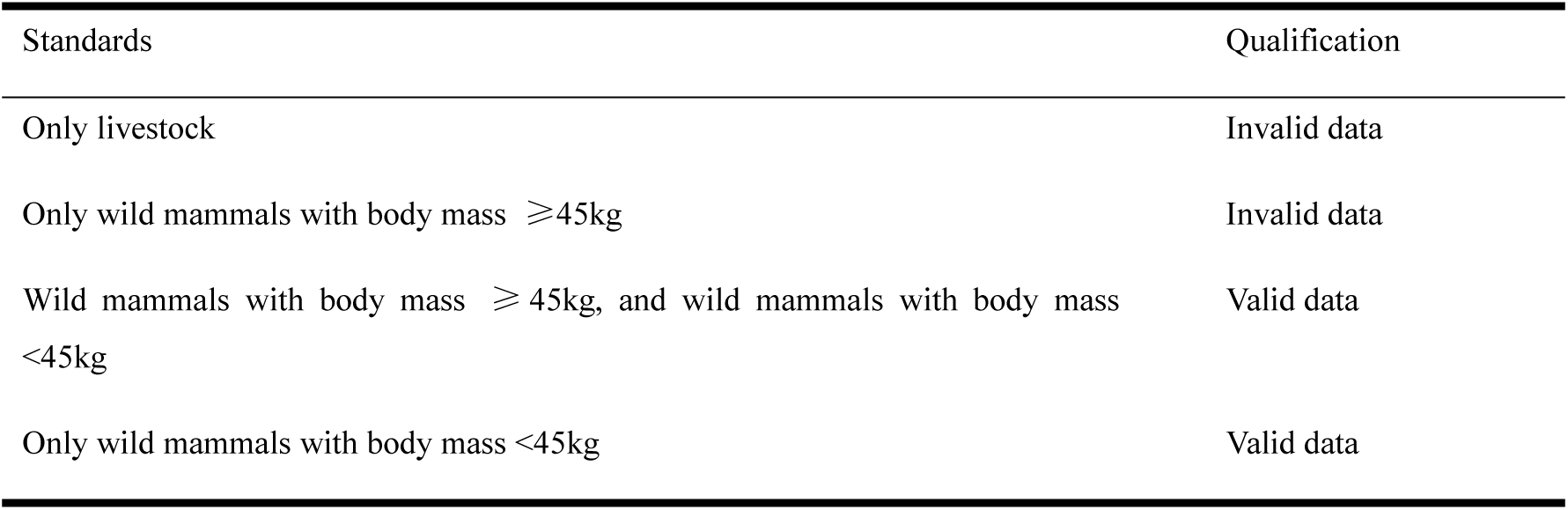
Qualifying the valid data.

**Table.S2.**
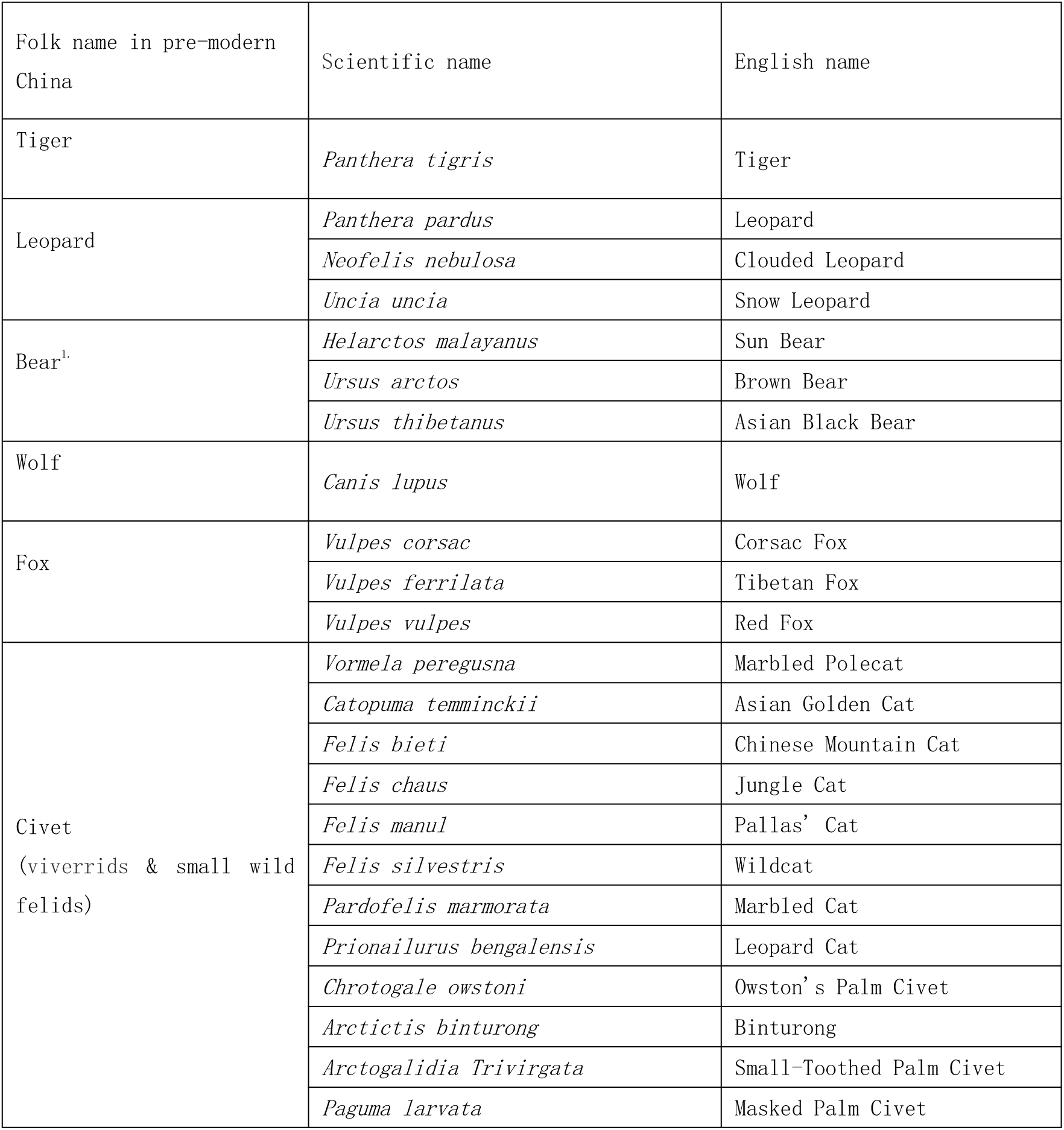
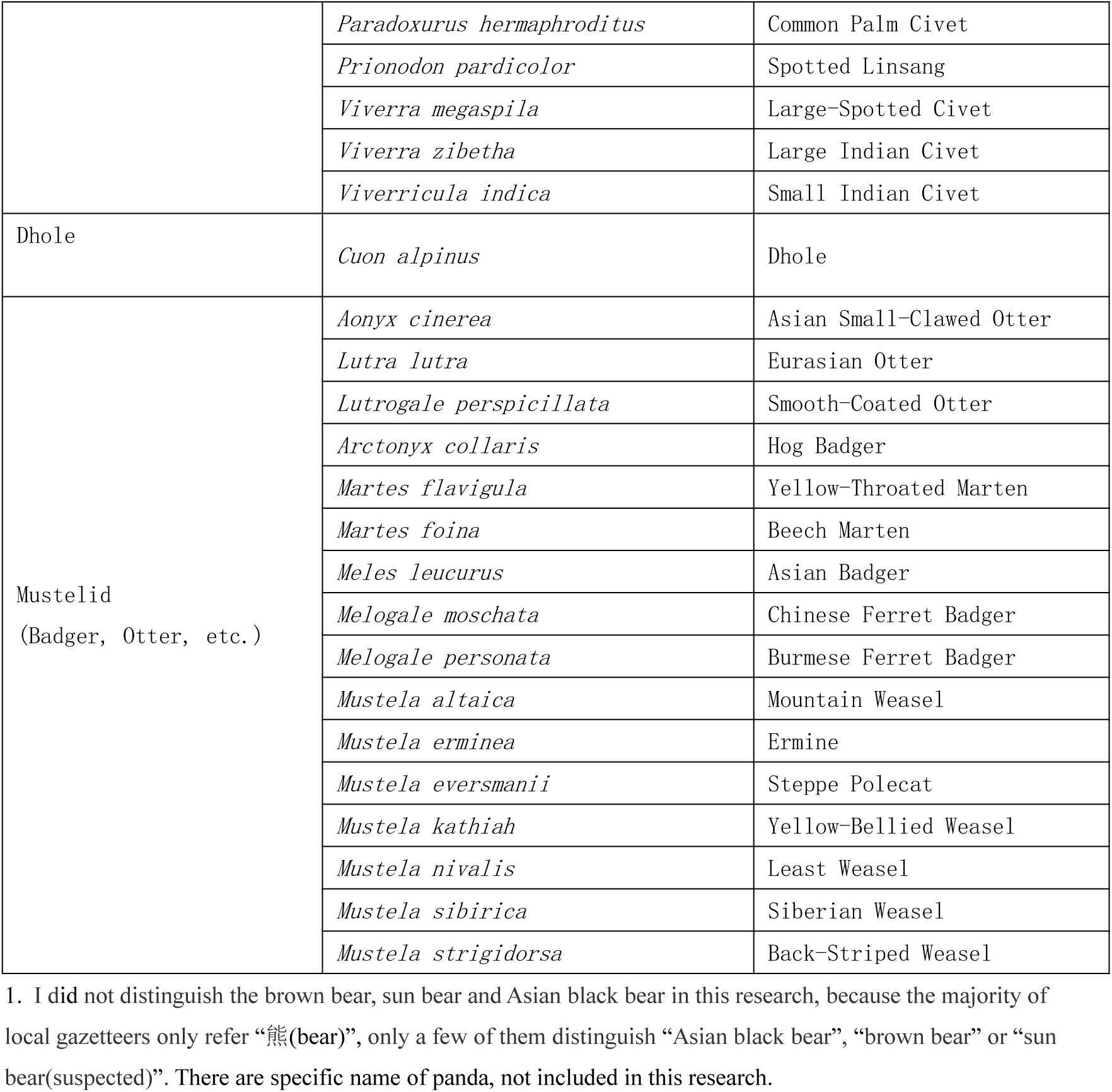
Folk name of predators in local gazetteers and relevant species in scientific taxonomy.

**Table.S3.** The change of average probability of carnivores from Period 1 to Period 4.

|  | Tigers | Leopards | Bears | Wolves | Foxes | Civets | Dholes | Mustelids |
| --- | --- | --- | --- | --- | --- | --- | --- | --- |
| Period 1 | 63.8% | 59.3% | 36.0% | 41.4% | 60.9% | 47.2% | 36.1% | 60.9% |
| Period 2 | 61.1% | 59.5% | 38.2% | 42.4% | 65.5% | 51.1% | 35.2% | 71.8% |
| Period 3 | 53.4% | 54.2% | 31.6% | 46.5% | 67.8% | 61.9% | 40.1% | 76.7% |
| Period 4 | 49.3% | 59.3% | 24.0% | 51.9% | 69.6% | 67.3% | 40.9% | 83.1% |

**Table.S4.** The average distribution probability in each altitude group in each period.

| alt1 |  |  |  |  |
| --- | --- | --- | --- | --- |
|  | P1 | P2 | P3 | P4 |
| Tigers | 37.2% | 29.6% | 28.4% | 21.5% |
| Leopards | 25.0% | 22.6% | 22.3% | 20.4% |
| Bears | 10.8% | 13.0% | 11.3% | 6.5% |
| Wolves | 45.6% | 43.4% | 43.8% | 33.3% |
| Foxes | 61.5% | 66.2% | 64.3% | 56.7% |
| Civets | 42.5% | 49.8% | 55.2% | 58.7% |
| Dholes | 16.4% | 18.0% | 17.2% | 17.7% |
| Mustelids | 72.7% | 86.1% | 87.5% | 80.0% |

| alt2 |  |  |  |  |
| --- | --- | --- | --- | --- |
|  | P1 | P2 | P3 | P4 |
| Tigers | 70.4% | 67.4% | 62.0% | 57.6% |
| Leopards | 59.5% | 61.2% | 54.3% | 56.4% |
| Bears | 31.0% | 32.8% | 26.8% | 16.7% |
| Wolves | 44.3% | 39.9% | 43.5% | 46.3% |
| Foxes | 59.8% | 62.7% | 60.0% | 62.0% |
| Civets | 53.3% | 64.4% | 73.8% | 74.5% |
| Dholes | 46.4% | 45.5% | 46.5% | 40.7% |
| Mustelids | 70.3% | 79.2% | 81.0% | 84.0% |

| alt3 |  |  |  |  |
| --- | --- | --- | --- | --- |
|  | P1 | P2 | P3 | P4 |
| Tigers | 79.9% | 72.3% | 64.1% | 59.2% |
| Leopards | 69.3% | 67.3% | 61.8% | 69.6% |
| Bears | 42.7% | 43.0% | 35.0% | 27.7% |
| Wolves | 34.7% | 33.9% | 37.9% | 48.7% |
| Foxes | 60.1% | 62.3% | 61.0% | 65.4% |
| Civets | 53.4% | 60.5% | 66.0% | 77.6% |
| Dholes | 46.4% | 44.6% | 44.7% | 45.6% |
| Mustelids | 63.7% | 73.0% | 75.6% | 83.1% |

| alt4 |  |  |  |  |
| --- | --- | --- | --- | --- |
|  | P1 | P2 | P3 | P4 |
| Tigers | 73.5% | 72.1% | 59.8% | 53.7% |
| Leopards | 70.0% | 69.8% | 64.1% | 70.6% |
| Bears | 45.4% | 45.2% | 34.8% | 27.0% |
| Wolves | 47.3% | 48.9% | 56.8% | 64.7% |
| Foxes | 66.8% | 68.7% | 73.3% | 77.7% |
| Civets | 47.9% | 47.5% | 57.6% | 65.3% |
| Dholes | 38.5% | 34.6% | 44.9% | 43.6% |
| Mustelids | 55.8% | 65.7% | 73.7% | 82.8% |

| alt5 |  |  |  |  |
| --- | --- | --- | --- | --- |
|  | P1 | P2 | P3 | P4 |
| Tigers | 57.9% | 64.2% | 52.8% | 54.4% |
| Leopards | 72.6% | 76.8% | 68.7% | 79.5% |
| Bears | 49.9% | 57.2% | 50.1% | 42.0% |
| Wolves | 35.2% | 46.2% | 50.5% | 66.7% |
| Foxes | 56.6% | 67.6% | 80.4% | 86.3% |
| Civets | 38.7% | 33.2% | 57.1% | 60.6% |
| Dholes | 32.9% | 33.4% | 47.4% | 57.1% |
| Mustelids | 41.9% | 55.0% | 65.4% | 85.7% |

**Table.S5.** Relations between each kind of carnivores and others (Kendall’s tau-b coefficient).

| Tigers |  |  |  |  |  |  |  |
| --- | --- | --- | --- | --- | --- | --- | --- |
|  | Leopards | Bears | Wolves | Foxes | Civets | Dholes | Mustelids |
| Period 1 | 0.551 | 0.410 | -0.076 | -0.009 | 0.288 | 0.554 | 0.139 |
| Period 2 | 0.613 | 0.396 | -0.047 | -0.079 | 0.191 | 0.480 | 0.068 |
| Period 3 | 0.613 | 0.400 | -0.046 | -0.070 | 0.240 | 0.452 | 0.113 |
| Period 4 | 0.603 | 0.479 | 0.017 | -0.172 | 0.232 | 0.481 | 0.054 |

| Leopards |  |  |  |  |  |  |  |
| --- | --- | --- | --- | --- | --- | --- | --- |
|  | Tigers | Bears | Wolves | Foxes | Civets | Dholes | Mustelids |
| Period 1 | 0.551 | 0.513 | -0.135 | -0.124 | 0.133 | 0.477 | -0.102 |
| Period 2 | 0.613 | 0.541 | -0.138 | -0.118 | 0.079 | 0.480 | -0.087 |
| Period 3 | 0.613 | 0.480 | -0.045 | 0.019 | 0.174 | 0.525 | 0.028 |
| Period 4 | 0.603 | 0.561 | 0.115 | 0.003 | 0.212 | 0.525 | 0.065 |

| Bears |  |  |  |  |  |  |  |
| --- | --- | --- | --- | --- | --- | --- | --- |
|  | Tigers | Leopards | Wolves | Foxes | Civets | Dholes | Mustelids |
| Period 1 | 0.410 | 0.513 | -0.125 | -0.096 | 0.166 | 0.418 | -0.075 |
| Period 2 | 0.396 | 0.541 | -0.117 | -0.146 | 0.001 | 0.381 | -0.128 |
| Period 3 | 0.400 | 0.480 | -0.088 | -0.041 | 0.132 | 0.382 | -0.039 |
| Period 4 | 0.479 | 0.561 | 0.030 | -0.139 | 0.197 | 0.385 | 0.030 |

| Wolves |  |  |  |  |  |  |  |
| --- | --- | --- | --- | --- | --- | --- | --- |
|  | Tigers | Leopards | Bears | Foxes | Civets | Dholes | Mustelids |
| Period 1 | -0.076 | -0.135 | -0.125 | 0.295 | -0.056 | -0.023 | 0.076 |
| Period 2 | -0.047 | -0.138 | -0.117 | 0.332 | -0.027 | -0.065 | 0.071 |
| Period 3 | -0.046 | -0.045 | -0.088 | 0.303 | -0.084 | -0.047 | 0.101 |
| Period 4 | 0.017 | 0.115 | 0.030 | 0.373 | -0.013 | 0.052 | 0.100 |
| Foxes |  |  |  |  |  |  |  |
|  | Tigers | Leopards | Bears | Wolves | Civets | Dholes | Mustelids |
| Period 1 | -0.009 | -0.124 | -0.096 | 0.295 | -0.044 | -0.034 | 0.151 |
| Period 2 | -0.079 | -0.118 | -0.146 | 0.332 | 0.017 | -0.132 | 0.121 |
| Period 3 | -0.070 | 0.019 | -0.041 | 0.303 | -0.025 | -0.014 | 0.163 |
| Period 4 | -0.172 | 0.003 | -0.139 | 0.373 | -0.059 | -0.042 | 0.208 |
| Civets |  |  |  |  |  |  |  |
|  | Tigers | Leopards | Bears | Wolves | Foxes | Dholes | Mustelids |
| Period 1 | 0.288 | 0.133 | 0.166 | -0.056 | -0.044 | 0.256 | 0.340 |
| Period 2 | 0.191 | 0.079 | 0.001 | -0.027 | 0.017 | 0.176 | 0.336 |
| Period 3 | 0.240 | 0.174 | 0.132 | -0.084 | -0.025 | 0.281 | 0.214 |
| Period 4 | 0.232 | 0.212 | 0.197 | -0.013 | -0.059 | 0.216 | 0.171 |
| Dholes |  |  |  |  |  |  |  |
|  | Tigers | Leopards | Bears | Wolves | Foxes | Civets | Mustelids |
| Period 1 | 0.554 | 0.477 | 0.418 | -0.023 | -0.034 | 0.256 | 0.124 |
| Period 2 | 0.480 | 0.480 | 0.381 | -0.065 | -0.132 | 0.176 | 0.056 |
| Period 3 | 0.452 | 0.525 | 0.382 | -0.047 | -0.014 | 0.281 | 0.048 |
| Period 4 | 0.481 | 0.525 | 0.385 | 0.052 | -0.042 | 0.216 | 0.092 |
| Mustelids |  |  |  |  |  |  |  |
|  | Tigers | Leopards | Bears | Wolves | Foxes | Civets | Dholes |
| Period 1 | 0.139 | -0.102 | -0.075 | 0.076 | 0.151 | 0.340 | 0.124 |
| Period 2 | 0.068 | -0.087 | -0.128 | 0.071 | 0.121 | 0.336 | 0.056 |
| Period 3 | 0.113 | 0.028 | -0.039 | 0.101 | 0.163 | 0.214 | 0.048 |
| Period 4 | 0.054 | 0.065 | 0.030 | 0.100 | 0.208 | 0.171 | 0.092 |

**Table.S6.** HPD Influence on carnivore distribution (Somers’ dyx)

|  | Tigers | Leopards | Bears | Wolves | Foxes | Civets | Dholes | Mustelids |
| --- | --- | --- | --- | --- | --- | --- | --- | --- |
| Period 1 | -0.112 | -0.296 | -0.293 | 0.030 | 0.042 | 0.028 | -0.190 | 0.181 |
| Period 2 | -0.161 | -0.304 | -0.351 | -0.053 | 0.013 | 0.150 | -0.129 | 0.215 |
| Period 3 | -0.134 | -0.264 | -0.290 | -0.100 | -0.113 | 0.054 | -0.188 | 0.179 |
| Period 4 | -0.254 | -0.381 | -0.289 | -0.294 | -0.173 | -0.059 | -0.222 | -0.034 |

**Table.S7.**
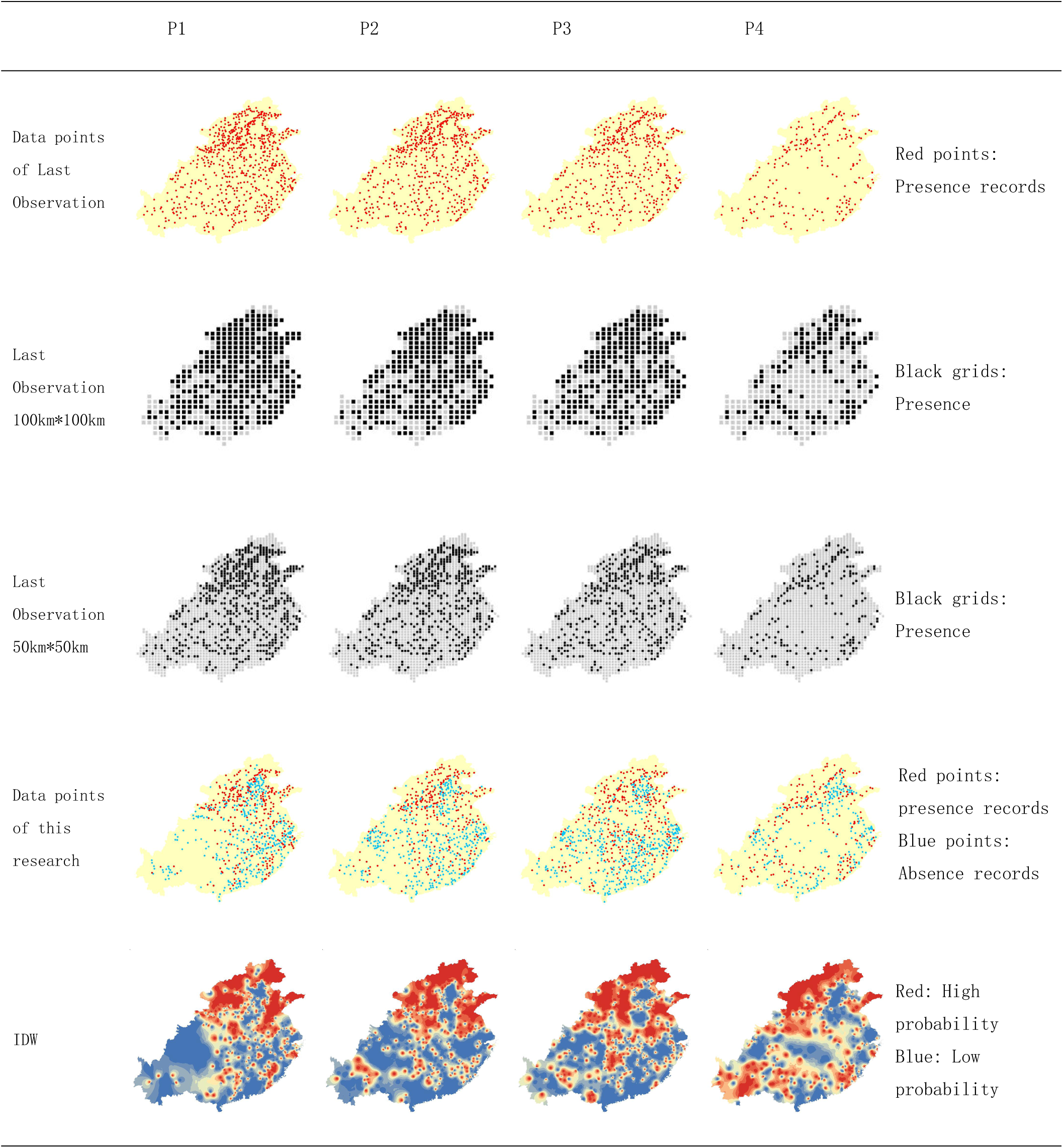
Criticizing the last observation precedure.

